# STEVE: Single-cell Transcriptomics Expression Visualization and Evaluation

**DOI:** 10.64898/2026.03.10.710898

**Authors:** Elijah Torbenson, Xiao Ma, Jhih-Rong Lin, Daniel J. Garry, Stephen C. Jameson, Zhengdong Zhang, Laura J. Niedernhofer, Lei Zhang, Meiyi Li, Xiao Dong

**Author notes:** Correspondence (M.L.), (X.D.).

## Abstract

Single-cell RNA sequencing (scRNA-seq) has become a key technology for characterizing cell-type heterogeneity in complex tissues. However, its utility depends on accurate and reproducible cell-type annotation, which remains a major analytical challenge. Although hundreds of computational tools have been developed for automated annotation, there is currently no systematic framework to evaluate annotation robustness in a dataset-specific manner or within the context of complete analytical pipelines. Here, we present STEVE (Single-cell Transcriptomics Expression Visualization and Evaluation), a quantitative framework designed to assess the accuracy, robustness, and reproducibility of cell-type annotation in scRNA-seq studies. STEVE implements three complementary in silico evaluation modules: (i) Subsampling Evaluation to quantify annotation stability under varying reference sizes and data partitions; (ii) Novel Cell Evaluation to assess the ability to detect previously unseen cell types; and (iii) Annotation Benchmarking to compare alternative annotation tools against ground-truth labels. In addition, STEVE includes a Reference Transfer Annotation module that enables cross-dataset cell-type mapping using external reference datasets. All modules are built upon a unified probabilistic framework that provides consistent confidence estimation across evaluation scenarios. We evaluated STEVE across four independent scRNA-seq datasets with experimentally defined or expert-curated cell-type labels. Our results show that annotation robustness is strongly influenced by the annotation method, biological separability, dataset complexity, and batch effects. STEVE provides a practical framework for quantifying annotation uncertainty and improving reproducibility in single-cell transcriptomic analyses. STEVE is freely available at GitHub (https://github.com/XiaoDongLab/STEVE).

## Introduction

Over the past decade, the rapid advancement of single-cell RNA sequencing (scRNA-seq) has made it an indispensable tool in developmental biology, immunology, cancer research, and aging studies. Its unique ability to preserve cell-specific gene expression profiles, often lost in bulk RNA sequencing, has greatly enhanced our understanding of cellular heterogeneity [1]. Despite its widespread adoption, scRNA-seq data analysis continues to face several technical challenges, particularly in the area of cell-type annotation [2]. The stability and resolution of upstream clustering critically influence annotation outcomes, as small perturbations in clustering parameters can substantially alter cell identity assignments, especially when cell types are transcriptionally similar or poorly separated [3]. Rare cell populations with low abundance are prone to being merged into larger, dominant clusters due to insufficient clustering resolution, leading to the masking of their distinct identities. In addition, conventionally, cell-type identification relied on expert-driven manual annotation, a time-consuming and often non-reproducible process across different research groups. This becomes a significant challenge, especially for large scale data integration project, e.g., the Human Cell Aging Transcriptome Atlas that we developed recently [4]. To address the challenge, automated annotation methods have been developed, broadly categorized into three methodological classes: marker gene-based identification (e.g., ScType [5], scCATCH [6], CellHint [7]), reference-based mapping (e.g., SingleR [8], scFseCluster [9], CellTypist [10], Cell BLAST [11]), and machine learning approaches (e.g., scDeepCluster [12]). As of now, more than 200 computational tools have been introduced to support these tasks [13].

Although the development of a wide array of cell-type annotation tools has enabled remarkable flexibility and innovation, it has also introduced the challenge of selecting the most suitable tool for a specific research question. To date, no benchmark study has comprehensively evaluated the performance of all 200-plus available annotation tools. Even among benchmark studies that have assessed smaller subsets, none have identified a tool that consistently outperforms others across all scenarios [14, 15]. As a result, it remains critically important to evaluate the accuracy and robustness of cell-type annotation on a study-specific basis. Given that annotation accuracy is influenced by multiple upstream analytical steps [16], a framework capable of evaluating annotation robustness within the context of a full pipeline is urgently needed. However, to our knowledge, no computational tool or pipeline currently exists to support this need.

To address this gap, we introduce *Single Cell Transcriptomics Expression Visualization and Evaluation* (STEVE), a novel evaluation framework for cell-type annotation in scRNA-seq experiments. STEVE is designed to operate on user-supplied datasets, optionally incorporating external reference datasets, and conducts three in silico experiments to cross-validate the annotation by different tools and/or to assess the reproducibility of a single tool’s output. As a proof of concept, we also implemented as part of STEVE a novel reference-based annotation method, which can be used independently if desired. STEVE offers a practical and accessible solution for researchers seeking to validate their cell-type annotations and improve the accuracy and reproducibility of scRNA-seq analyses.

## Materials and Methods

### Public scRNA-seq datasets

To demonstrate the usage and performance of STEVE, we applied STEVE to several scRNA-seq datasets available from the literature. Stewart et al [17] experimentally separated B lymphocytes from a healthy male volunteer with fluorescence-activated cell sorting (FACS) into five groups: IgM Memory Cells (CD19+IgD+CD27+), Double Negative cells (CD19+IgD-CD27-), Classical Memory cells (CD19+IgD-CD27+), Naïve cells, (CD19+IgD+CD27-CD10-), and Transitional cells (CD19+IgD+CD27-CD10+). Each group was then sequenced separately, thus allowing for a ground-truth annotation of cell type experimentally. The PBMC data from the Tabula Sapiens Consortium was also used to provide ground truth cell-type classification [18]. While these cells were not separated experimentally, they were annotated by a large collaboration of experts. Peripheral blood mononuclear cells (PBMCs) isolated from a healthy male donor from 10X genomics were also used as a dataset (5k Human PBMCs Stained with TotalSeqTM-C Human TBNK Cocktail, Chromium NextGEM Single Cell 5’ [WWW Document], n.d. 10× Genomics. URL https://www.10xgenomics.com/datasets/5k-human-pbmcs-stained-with-totalseq-C-human-TBNK-cocktail-NextGEM (accessed 5.23.25)) to evaluate the performance of STEVE. Finally, single-nucleus RNA sequencing (snRNA-seq) data generated from mouse cardiomyocytes in the Cui et al. study in 2021 was also used [19]. From the above, we used their expression level matrices and performed data cleaning using Seurat R package [20].

### STEVE

The accuracy and robustness of cell-type annotation depend not only on cell-type-specific features, whether derived from annotation tools or human expertise, but also heavily on the quality of upstream cell clustering. To evaluate these factors, STEVE implements three distinct in silico experiments, i.e., *Subsampling Evaluation, Novel Cell Evaluation*, and *Annotation Benchmarking*, each targeting a different source of variability in annotation performance. These experiments can be run independently, allowing users to tailor the evaluation to their specific study design.

#### *STEVE model* (Figure 1A)

The computational core of STEVE is a cell-cluster (or “type”) mapping method, which are used in the three in silico experiments. Specifically, to assign cell types to query cells, we implemented a density-based Bayesian classification framework in UMAP space (the dimensional reduction method can be changed as needed). First, after integration and dimensionality reduction, we embedded a reference (ground truth) and query (user) cells into a shared two-dimensional UMAP (or tSNE depending on the user’s needs) space. For each reference cell type *c*, we modeled its spatial distribution in UMAP space using two-dimensional kernel density estimation (KDE) implemented via kde2d (MASS package) [21]. This yielded a smoothed density surface

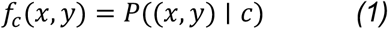

representing the likelihood of observing a cell at location (*x,y*) given cell type *c*. To enable probabilistic inference, each density surface was discretized on a fixed grid and normalized so that the total density across the grid summed to one. This produced a discrete likelihood function for each cell type.

We then estimated cell-type priors *P*(*c*) from the reference dataset using empirical frequencies with Laplace smoothing. For each grid location, posterior probabilities were computed via Bayes’ theorem:

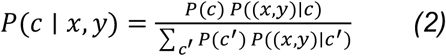

Thus, each position in UMAP space was associated with a posterior probability vector across all reference cell types.

For each query cell, its UMAP coordinates were mapped to the nearest grid location, and posterior probabilities were extracted. Cell-type assignment was determined by the maximum a posteriori estimate. To control classification confidence, we computed the posterior odds ratio between the most probable and second-most probable cell types:

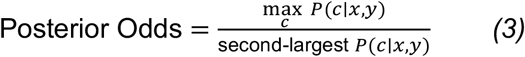

A cell was assigned to the top-ranked cell type only if the posterior odds exceeded a predefined threshold (default = 2). Otherwise, the cell was labeled as unassigned, reflecting insufficient separation in posterior probability and potential overlap between cell-type density distributions. This framework integrates spatial density modeling and Bayesian inference to quantify both assignment probability and classification confidence for each individual cell.

#### Subsampling Evaluation (Figure 1B)

STEVE evaluates the robustness of clustering and its impact on annotation accuracy through a series of reference subsampling experiments. In each iteration, a dataset is split into unequal partitions (e.g., 10% vs. 90%, 20% vs. 80%, …, up to 90% vs. 10%), with the first subset used as the reference to annotate the other subset (denoted as user dataset). Sensitivity and specificity are computed for each split to quantify the robustness of annotation under limited reference data conditions.

#### Novel Cell Evaluation (Figure 1B)

STEVE assesses the pipeline’s ability to correctly identify novel or unseen cell types using a novel cell detection framework. A dataset is split evenly into a reference and a user set, and in each iteration, a specific cell type is removed from the reference. The expectation is that cells of this omitted type in the user set should be labeled as “unknown” or “novel” when annotated against the reduced reference. STEVE reports both the correct identification rate of these novel cells and the frequency of misclassification into known types.

#### Annotation Benchmarking (Figure 1B)

STEVE enables annotation benchmarking when ground-truth labels are available, allowing for evaluation of annotation methods, including automated tools and manual expert annotations. The dataset is partitioned into reference and user subsets, with the user set annotated using the method to be evaluated. STEVE then compares these user-supplied labels to the ground truth using sensitivity, specificity, and novel cell detection metrics, offering a standardized way to benchmark annotation accuracy.

Finally, the STEVE model can be directly used to annotate a user-supplied dataset based on a second, external dataset with known ground-truth labels. We denoted this module as *Reference Transfer Annotation*. This functionality enables STEVE to act not only as an evaluation framework but also as a practical annotation tool for new datasets, leveraging high-quality reference data from previous studies.

## Results

### Subsampling Evaluation

We applied the *Subsampling Evaluation* module of STEVE to four published scRNA-seq datasets: Stewart et al., blood of Tabula Sapiens, 10x Genomics PBMCs, and Cui et al. (See **Materials and Methods**). Each dataset included both the single-cell gene expression count matrix and accompanying cell-type annotations as defined in the original publications. For each dataset, cells were randomly split into two subsets: a reference set and a user set. The reference set was used to annotate the user set, and the predicted annotations were then compared to the original labels to assess accuracy. This process was repeated iteratively using varying reference-to-user split ratios, ranging from 1:9, 2:8, …, to 9:1. In principle, since the reference and user cells are randomly sampled from the same dataset, the predicted annotations should closely match the original annotations. Any loss in annotation accuracy, quantified by sensitivity and specificity, can thus be attributed to reduced robustness in cell clustering as well as its upstream pipeline steps, e.g., feature selection and normalization.

The results across all four datasets support this rationale. The Stewart et al. (**Figures 2A–C, S1 and S2**) and 10x PBMCs (**Figures 2D–F and S3**) datasets achieved the highest average sensitivity (88% and 97%, respectively) and specificity (97% and 100%, respectively), likely reflecting their high data quality and well-controlled experimental conditions. In contrast, the blood of Tabula Sapiens dataset (**Figures 2G–I and S4**) exhibited lower sensitivity (average 62%), although it maintained high specificity (average 99%). This discrepancy may be due to the fact that nearly 85% of the cells are from 4 cell types and the rest of the 15% are from 14 cell types. It is likely harder to correctly annotate these smaller cell populations. The dataset from Cui et al. (**Figures 2J–L and S5**) showed the lowest average sensitivity and specificity (52% and 87%, respectively). This likely reflects the inherent challenges in annotating cardiomyocyte subtypes, which are less well-characterized compared to immune cell subsets. While B cell subtypes (e.g., in the Stewart dataset) are well-defined and supported by canonical markers, cardiomyocyte subtypes remain ambiguous in the literature, commonly grouped into five types (CM1–CM5) [19], despite clear evidence of more diverse subpopulations in the UMAP embeddings (**Figures 2J–K**). Of note, different split ratio does not have a substantial impact on sensitivity or specificity scores in any of the above datasets, suggesting the number of cells in the experiments are sufficient for cell clustering and annotation. Overall, these results illustrate that the resolution and robustness of cell-type annotation are highly context dependent and influenced by both biological and technical factors.

**Figure 1.**
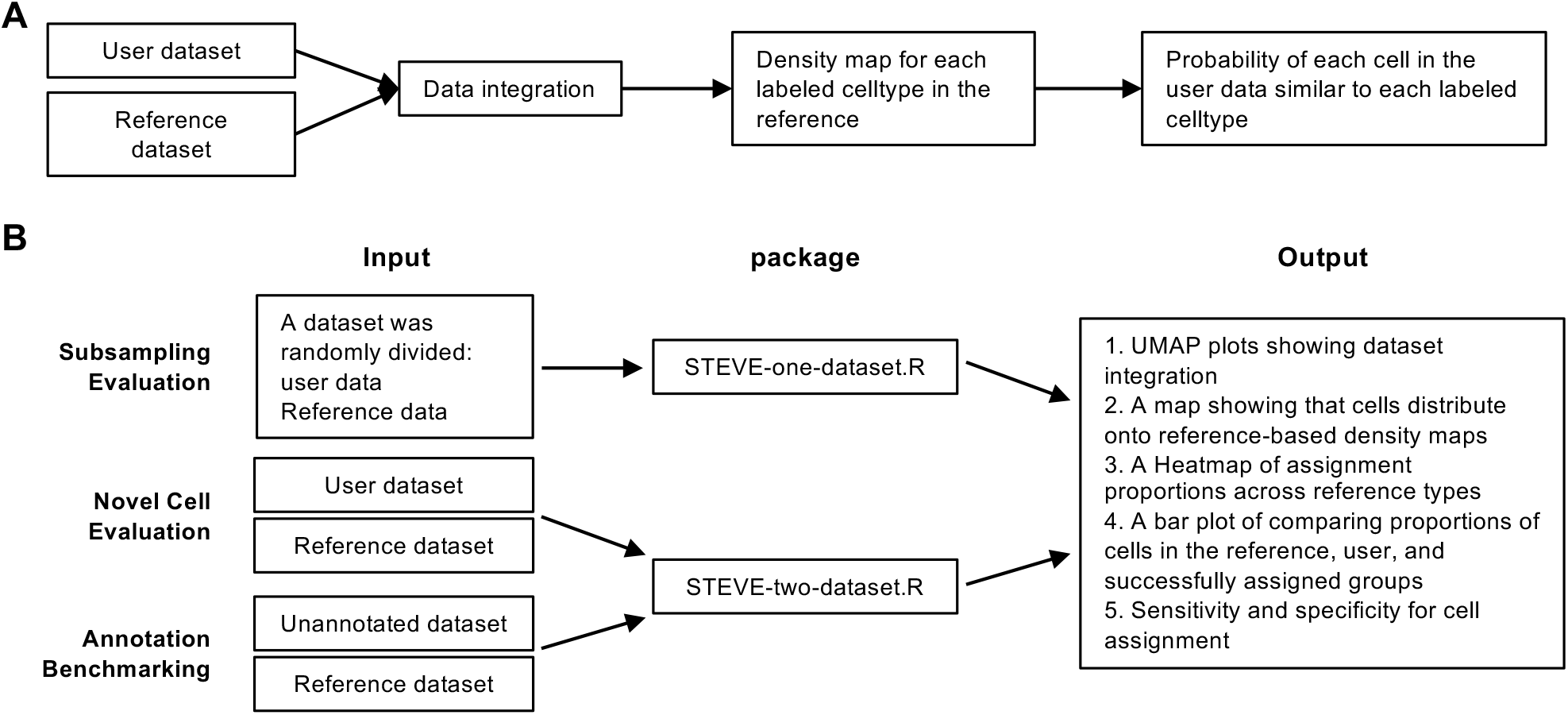
Overview of STEVE model. *(A)* A schematic overview of STEVE model. *(B) Subsampling Evaluation:* To compare two different annotation methods on the same dataset, randomly divide the dataset into two halves, one designated as the user data and the other as the reference data, then execute the script STEVE-one-dataset.R which will automatically repeat the annotation algorithm ten times. *Novel Cell Evaluation:* To compare annotations between two distinct datasets, assign one dataset as the user data and the other as the reference data, then run the script STEVE-two-datasets.R. *Annotation Benchmarking:* To annotate a dataset using a reference dataset, assign the target dataset as the user data and ensure that the celltype_groundtruth column is set to “X”, then execute the script STEVE-two-datasets.R. See our GitHub link for detailed usage instructions.

**Figure 2.**
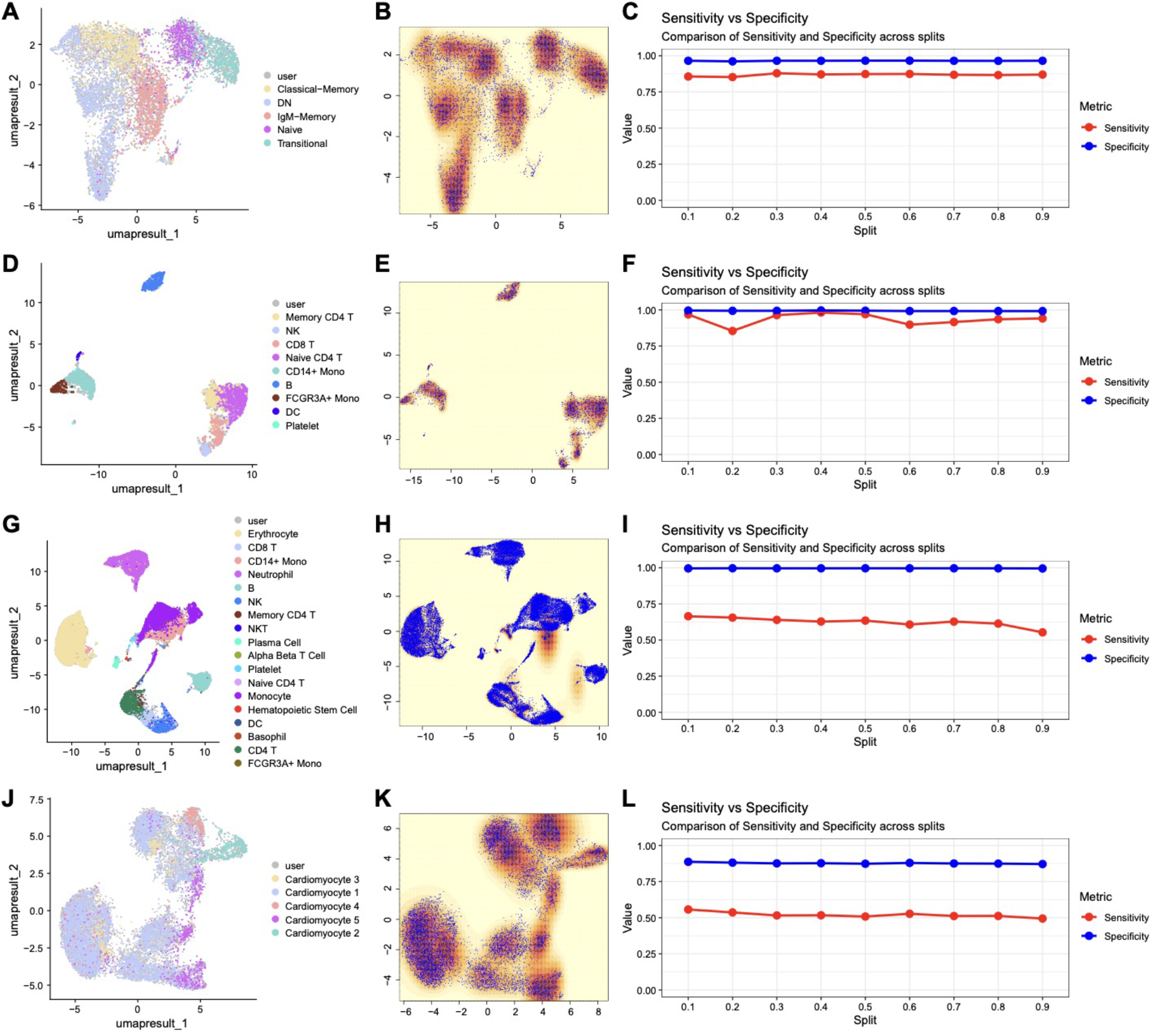
Subsampling Evaluation. We split the Stewart et al. dataset into unequal fractions (10% vs 90%), with the 10% being used to annotate the 90% and compute a sensitivity and specificity score. In the next iteration, the dataset was split into 20% vs 80%, with the 20% being used to annotate the 80% and compute a sensitivity and specificity score (and so on). Figure 2A shows a UMAP visualization of the entire Stewart et al. dataset, with the reference portion being labeled by cell type and the rest designated as the user dataset for subsequent benchmarking analysis. Figure 2B shows the density map under 50% vs 50% split. Figure 2C shows the sensitivity and specificity measurements across different split ratios.Figures 2D-F, 2G-I, and 2J-L were generated using the same pattern on the 10X dataset, Tabula Sapiens dataset, and Cui et al. dataset.

### Novel Cell Evaluation

The *Novel Cell Evaluation* module in STEVE enables users to assess whether previously unseen or novel cell types can be correctly detected when mapping cells against a predefined reference. As shown in the Subsampling Evaluation, the 1:1 split ratio has minimal impact on annotation accuracy in these datasets. In each iteration of the *Novel Cell Evaluation*, one cell type was removed from the reference set. The remaining reference cells were then used to annotate the full user subset. Cells belonging to the omitted type in the user set were expected to be classified as “unknown” or “uncertain,” rather than incorrectly assigned to one of the remaining reference types. With the above, *Novel Cell Evaluation* challenges the clustering process to detect and appropriately handle cell types absent from the reference, a scenario common in exploratory or cross-tissue studies.

We applied this module to the Stewart (**Figures 3A-D**) and Cui (**Figures 3E-H**) datasets, which include both gene expression matrices and detailed cell-type annotations. The other two datasets, 10x Genomics PBMCs (**Figure S6**) and blood of Tabula Sapiens (**Figure S7**), were also used in this experiment. The Stewart dataset has an average sensitivity of 40% and specificity of 100%, whereas Cui et al. had an average sensitivity of 16% and specificity of 100%. In the Stewart dataset, most classical memory, double-negative, and IgM-memory B cells were frequently misclassified as one another due to their similar characteristics. Naïve and transitional B cells were also often predicted as each other. In mouse cardiomyocytes, types 2– 5 were frequently predicted as type 1, likely because their defining features are less specific and substantially overlap with those of type 1. The tabula Sapiens dataset had an average sensitivity of 27% and specificity of 100%, with certain cell types performing much better than others, such as the CD4 T, NK T cell, Plasma cell, and Platelet cell having sensitivities over 60% (**Figure S7**). The 10X Genomics dataset performed poorly, having an average sensitivity of 2% and specificity of 100% (**Figure S6**). Overall, the results suggest that the more distinctly a cell type is characterized and the larger its proportion within a dataset, the more likely successful identification as a novel cell type. Conversely, when a cell type is predicted to be novel, the prediction is typically correct, as reflected by the consistently high specificity. These plots also highlight potential mismatches between cell-type frequency and classification performance, with some rare cell types exhibiting disproportionately high classification success.

**Figure 3.**
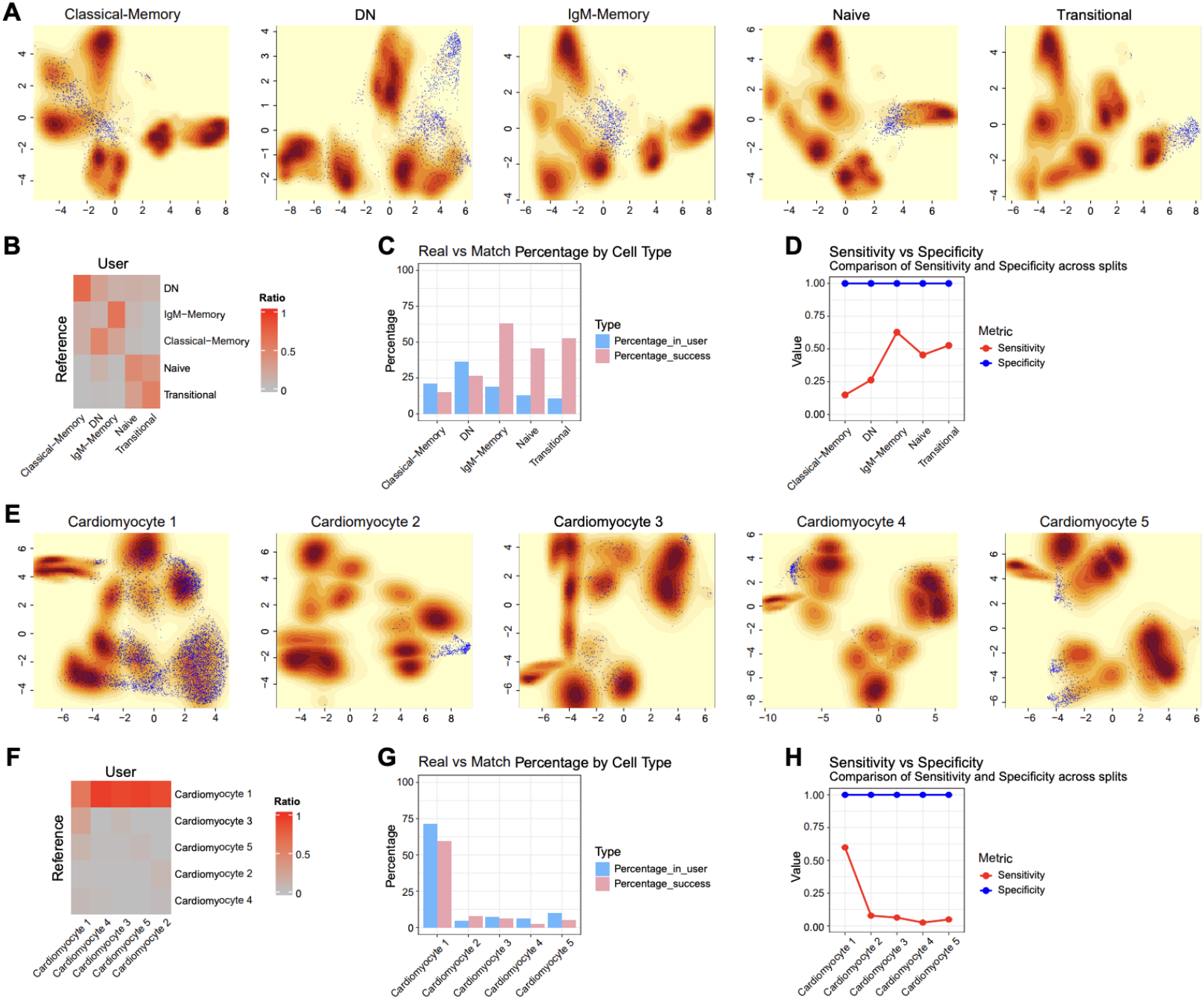
Novel Cell Evaluation. We split the dataset and used one portion as a reference to annotate the other portion, but this time we removed a cell type from the reference portion. In principle, the cells from the same cell type in the user should thus be annotated as “uncertain” or “new.” The first image in Figure 3A shows a density map embedded for the reference portion of the dataset without the contribution of Classical-Memory cells. For visualization purposes, the Classical-Memory cells from the user half are overlaid onto the density map to show the lack of overlap with the reference-density map. The remaining images of Figure 3A follow the same pattern, with the only difference being the DN, IgM-Memory, Naïve, and Transitional cells were removed from the reference. Figure 3B is a misclassification matrix that shows cell types that the cells were incorrectly assigned to, with the “unknown/new” label from the user part (shown as the column names) being replaced with the name of the corresponding cell from the reference part (shown as the row names) that was removed. Figure 3C shows both the percentage of total user cells made up of new/unknown cells, as well as the percentage of those new/unknown cells correctly classified as “new/unknown.” Figure 3D shows the sensitivity and specificity for each iteration where a cell type was removed from the reference. Figure 3E-H follow the same pattern as Figure3A-D, but this time for the Cui et al. dataset.

### Annotation Benchmarking

To demonstrate STEVE’s function to evaluate performance of annotation tools based on dataset with known ground-truth annotations, we performed *Annotation Benchmarking* of two popular cell type annotation tools, scType [5] and singleR [8]. To include potential impact of cell clustering, the *Annotation Benchmarking* module also randomly splits a dataset into two (of equal sizes). An annotation tool, e.g., singleR, is applied to one half and the resulted annotation is compared to the ground-truth annotation of the other half after data integration.

We tested this approach on the 10x Genomics PBMC dataset (**Figure 4**), which contains relatively few cell types with more balanced proportions. In this case, scType outperformed singleR: while both tools achieved 100% specificity, scType exhibited substantially higher sensitivity across most cell types.

**Figure 4.**
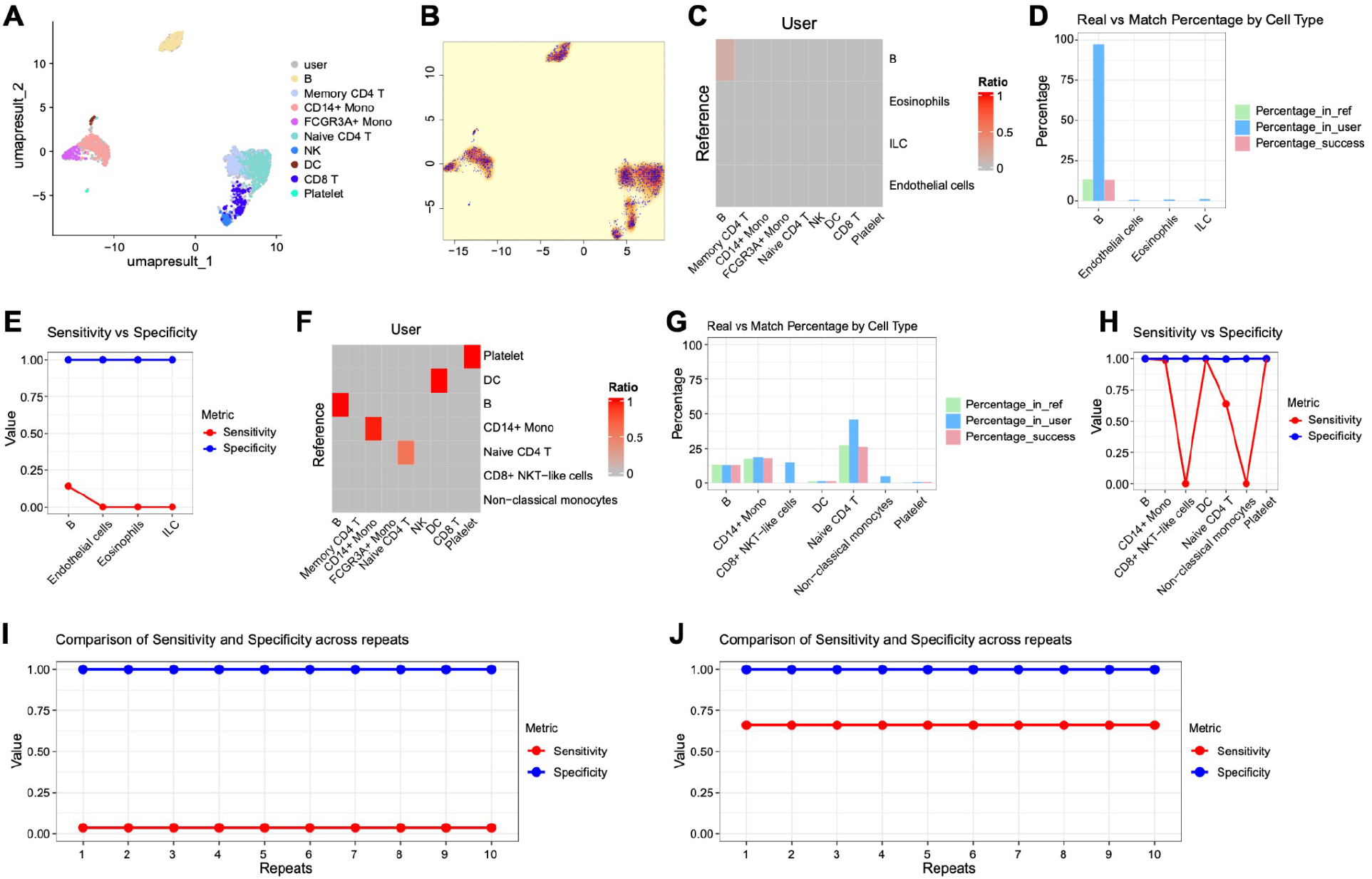
Annotation Benchmarking. In this set of experiments, the 10X dataset was split into a user half and a reference half, with both scType and singleR being used as the annotation tool. Figure 4A-B just show a cell-labeled and density-map UMAP of the dataset for visualization (nothing to do with singleR or scType, purely for visualization purposes). scType and singleR were then each used to annotate the 10X dataset. The resulting annotations were then considered “ground truth.” The 10X dataset was then split into half, with the reference half (made available by the earlier ground truth annotations) to annotate the other half across multiple repeats. Figures 5C-E and I are from singleR’s performance evaluated by STEVE, and Figures 5 F-H and J are from scType’s performance evaluated by STEVE.

### Reference Transfer Annotation

Finally, we included in STEVE a *Reference Transfer Annotation* module, which enables users to annotate cell types of their own dataset based on the ground-truth annotation of a reference dataset. This is achieved by integrating two datasets together in a combined dataset [22], and the grand-truth annotation is “transferred” to user’s dataset based on the density distribution of the corresponding cell types of the reference data. **Figure 5** shows examples of this process with the datasets that we collected above. The 10X dataset was first used as reference to annotate the Stewart dataset, with resultant 99% sensitivity and 100% specificity (**Figures 5A-E**). Next, the Tabula sapiens dataset was used to annotate the Stewart dataset, with resultant 99% sensitivity and 100% specificity. Finally, the Tabula sapiens dataset was used to annotate the 10X dataset, with 0, 0, 81%, 0, 98%, 5%, 5%, 100%, 86% sensitivity and 100%, 100%, 97%, 100%, 100%, 99%, 95%, 97%, 100% specificity for the memory CD4 T cells, Naive CD4 T, CD8 T, DC, B cells, CD14+ Monocyte cells, FCGR3A+ Monocyte cells, NK, Platelet. Overall, the *Reference Transfer Annotation* module offers a light-weight reference-based annotation tool in addition to the evaluations above.

**Figure 5.**
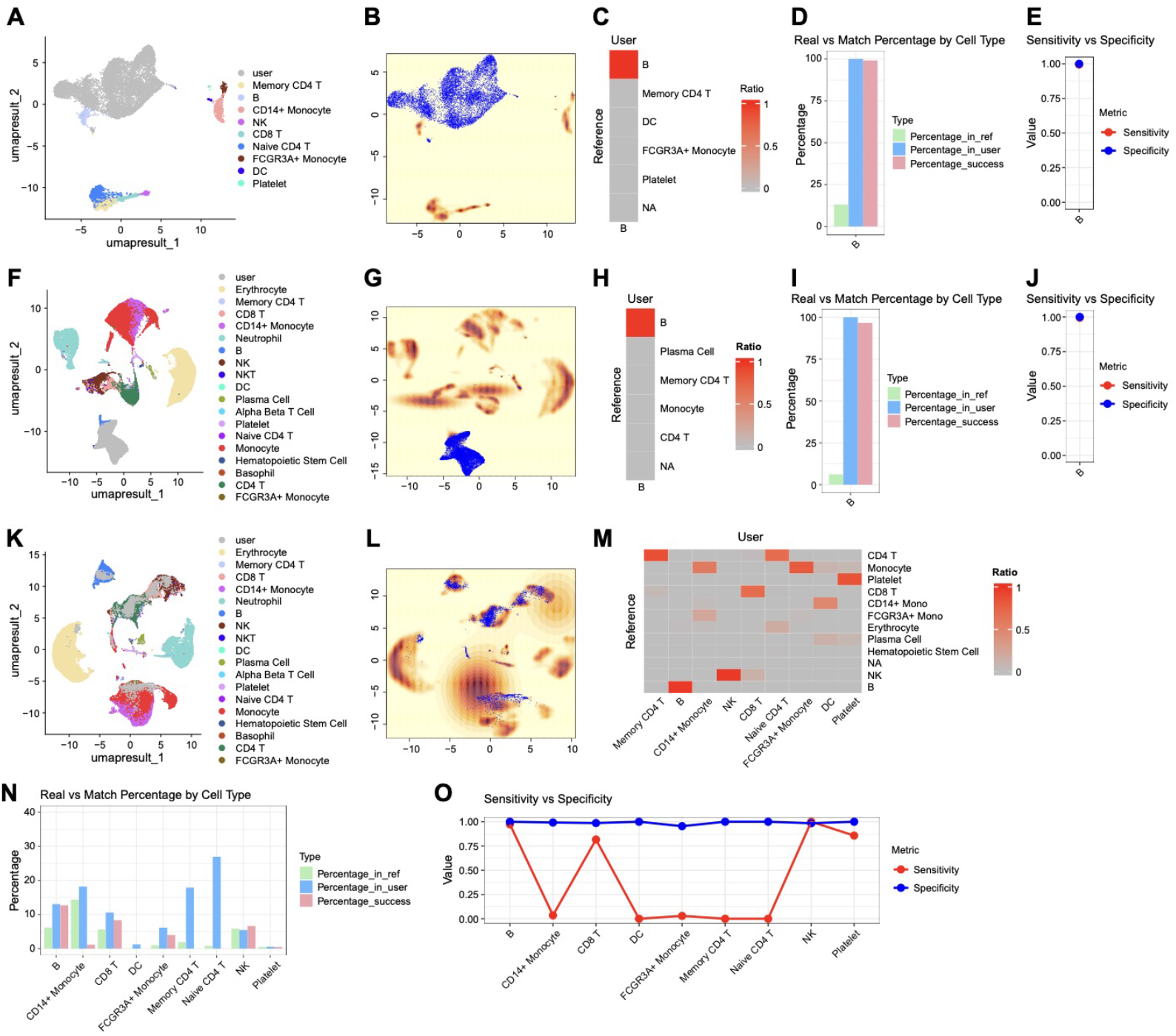
Reference Transfer Annotation. In this next experiment we used one dataset as reference to annotate another dataset (as opposed to splitting one in half) To generate Figures 5A-E, the 10X dataset was used to annotate the Stewart dataset. Note that the Stewart dataset is composed of B cell subtypes whereas 10X only has the general category of B cell, so all the subtypes in Stewart were relabeled as just “B cell”. To generate Figures 5F-J, the Tabula Sapiens dataset was used to annotate the Stewart et al. dataset. To generate Figures 5K-O, the tabula sapiens dataset was used to annotate the 10X dataset.

## Discussion

We present STEVE, a bioinformatic tool to evaluate the robustness and quantify the uncertainty of the cell-type annotation process with the use of three in silico experiments (or modules). We validated the effectiveness of the first module, *Subsampling Evaluation*, by applying it to four published scRNA-seq datasets. Because decreases in performance metrics generated by the *Subsampling Evaluation* module can be attributed to either the technical noise inherent in upstream pipeline steps or difficult-to-characterize biological variability in the dataset itself, we expected that datasets with excellently-controlled experimental conditions and well-demarcated cell types would score highest in this first module, and this is what we saw. The Stewart et al. dataset, which was generated by gold-standard FACS, had the highest sensitivity and specificity, whereas the Tabula Sapiens dataset, which is a conglomeration of data from many different institutions and likely biased with data integration issues, had poorer performance. The second module, *Novel Cell Evaluation*, gauges the ability of the user’s pipeline to identify novel cell types, and we validated this module by applying it to datasets with biologically easy-to-distinguish cell types versus harder-to-distinguish cell types (e.g. cell subtypes). As expected, the datasets with limited biological heterogeneity (e.g. the cardiomyocyte subtypes in the Cui et al. dataset) had lower performance metrics than datasets with well-demarcated cell types (e.g. Tabula Sapiens dataset). The third module, *Annotation Benchmarking*, allows the user to directly compare the performance of other published cell-annotation tools, and we demonstrated this by comparing the performance of singleR and scType. Each tool was used to annotate the Stewart et al. dataset, and these annotations were used in place of the ground-truth annotations, with the dataset being subsequently subjected to a similar evaluation process as in module 1, with the overall performance thus reflecting upon the validity of the annotation tools used beforehand. We found that while exhibiting similar specificities, scType had better sensitivity than singleR. Finally, we also include a fourth module called *Reference Transfer Annotation*. While this module is not used for evaluation of the cell-annotation process, it can be used as an annotation tool itself in lieu of other published tools such as scType and singleR.

Rather than benchmarking the performance of popularly-used tools at one specific step of the pipeline (e.g. normalization or cell annotation), STEVE serves as a quality control engine for entire pipelines by using the performance of the cell-annotation step as an integrative indicator of overall pipeline integrity. While not being a benchmarking tool per se, we found that our conclusions generated by STEVE generally matches what has been found in scRNA-seq benchmarking studies. For instance, performance was best when annotating majorly different cell types (e.g. B cells vs T cells) and lower when annotating very similar cell types and subtypes (e.g. cardiomyocytes), a finding found in previous benchmarking studies [23, 24]. Another benchmarking study found that while reference datasets can have multiple levels of bias (individual level, condition level, and batch level), the higher-level experimental batch effect biases contributed the most to decreased performance [25, 26]. Our study reproduced this conclusion, as the Tabula Sapiens dataset was the most subject to batch effect biases and performed worse than single-experiment datasets, such as the Stewart et al. dataset. The same benchmarking study also found that increasing dataset complexity (i.e. increased number of cell types) led to more difficult annotation, a finding also reproduced by STEVE: the Stewart et al. dataset and 10X dataset, both with less than 10 cell types, performed better in the *Subsampling Evaluation* module than Tabula Sapiens which had almost double the cell types. The exception to this was the Cui et al. dataset which had only four cell types but performed the worse, likely due to the insufficient of transcriptomic variation between the cardiomyocyte subtypes.

We envision that researchers can use STEVE as both a quality assurance tool for their data pipeline and as a method to select the proper cell annotation tool: With over 200 single cell-annotation tools available, it can be difficult to select the proper tool to use for analysis. Using STEVE, researchers can upload their own dataset and compare the performance of various annotation tools in the context of their local environment, similar to how we compared singleR and scType. In addition, no data pipeline is free of bias, and variations in pipeline often shift where the bias lies rather than getting rid of it all together. With using STEVE, researchers are able to make informed decisions of where their bias may be: for instance, exchanging an upstream normalization algorithm with another may lead to increased performance in the *Subsampling Evaluation* module, but decreased performance in the *Novel Cell Evaluation* module, thus alerting researchers that their current pipeline may have increased accuracy in annotating known cell types but increased difficulty with identifying new cell types. The researchers can thus adjust their pipeline according to their priorities. Additionally, the fourth module *Reference Annotation Transfer* of STEVE does serve as a valid cell-annotation tool independent of evaluation, and researchers are encouraged to try this in their pipeline.

Notably, many of the biases discussed above, such as excessive numbers of cell types, insufficiently demarcated cell populations, or substantial batch effects, substantially limit the robustness and accuracy of cell-type annotation, as demonstrated by STEVE. These challenges likely arise in large part from the inherent sparsity and dropout characteristic of most single-cell RNA-seq protocols [27], which constrain the resolution at which subtle cell subtypes or dynamic cellular states (e.g., changes during aging [28]) can be reliably detected. By applying STEVE to a given dataset, researchers can systematically probe the upper bound of achievable annotation accuracy imposed by the data itself, thereby distinguishing limitations intrinsic to experimental design and data quality from those introduced by downstream computational pipelines.

As with all tools, STEVE does have some limitations. The most important one is that the veracity of the performance metrics in the modules depend upon the accuracy of the pre-existing cell type annotations of the reference dataset. Accordingly, the cell type labels in the reference dataset should ideally come from experimental means (e.g. FACS). If this is not possible, the annotations should come from methodology not dependent upon automated cell annotation tools (e.g. expert annotation, as used in the Tabula Sapiens dataset).

Overall, we developed STEVE to evaluate the robustness and accuracy of cell-type annotation, which is one of the most critical steps and components in single-cell RNA sequencing experiments. In the future, we hope to expand the evaluation to other steps, e.g., normalization, batch-effect correction, and downstream analysis, e.g., trajectory analysis [29] and cell-cell communication [30], as well as the analysis of spatial transcriptomics [31], to enhance the reproducibility and scientific rigor of the omics experiment at the single-cell level.

## Code availability

The source code and usage instructions of STEVE is freely available at https://github.com/XiaoDongLab/STEVE.

## Acknowledgements

This work was supported by the U.S. National Institutes of Health (U19 AG056278, P01 AI172501, P01 HL160476, and U54 AG076041). The funders had no role in study design, data collection and analysis, decision to publish, or preparation of the manuscript.

