## Supplemental Figures 1-7 for "STEVE: Single-cell Transcriptomics Expression Visualization and Evaluation"

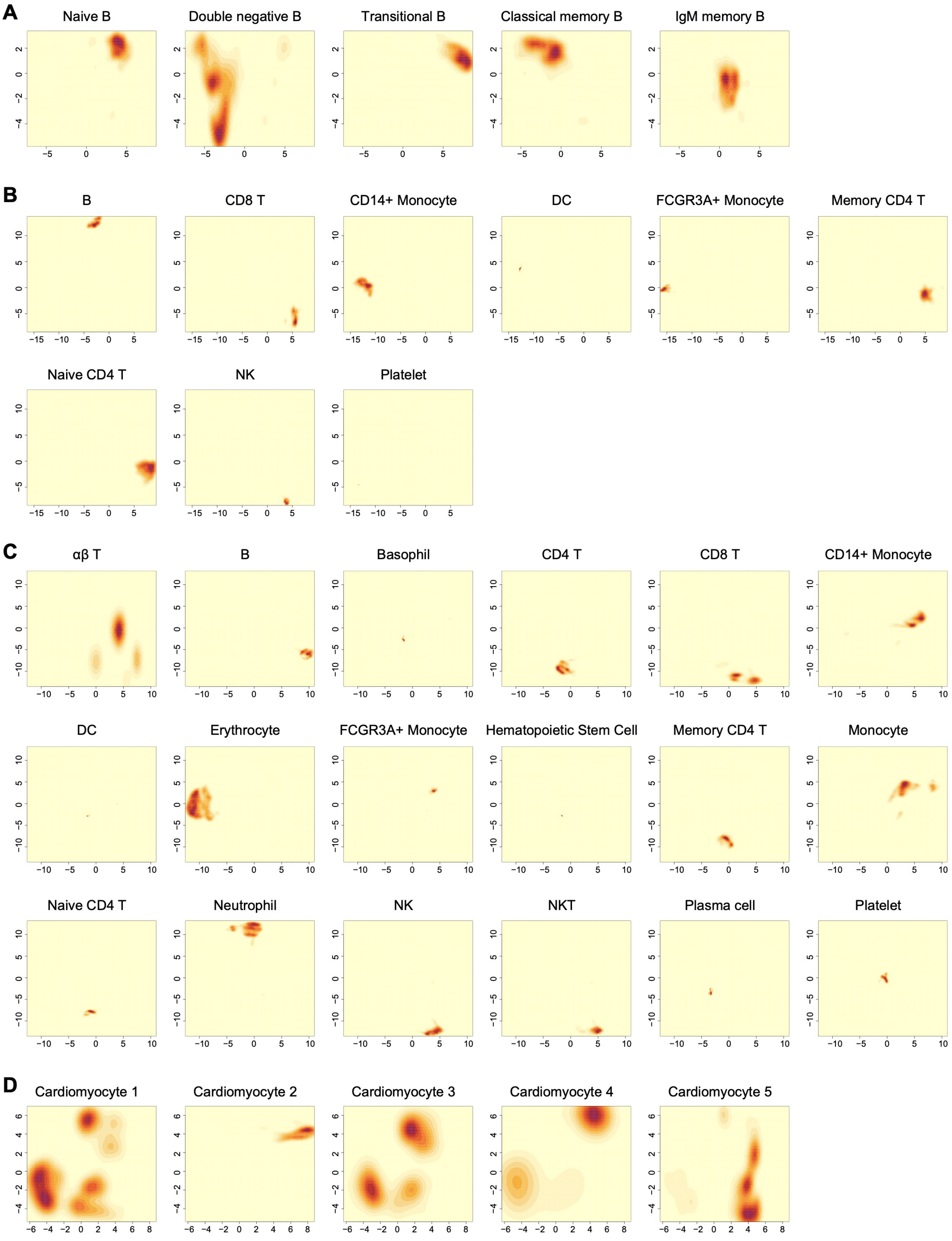


***Figure S1. Density maps based on the reference dataset's labeled cell types.***

*Individually supporting Figures 2b, 2e, 2h, and 2k)*

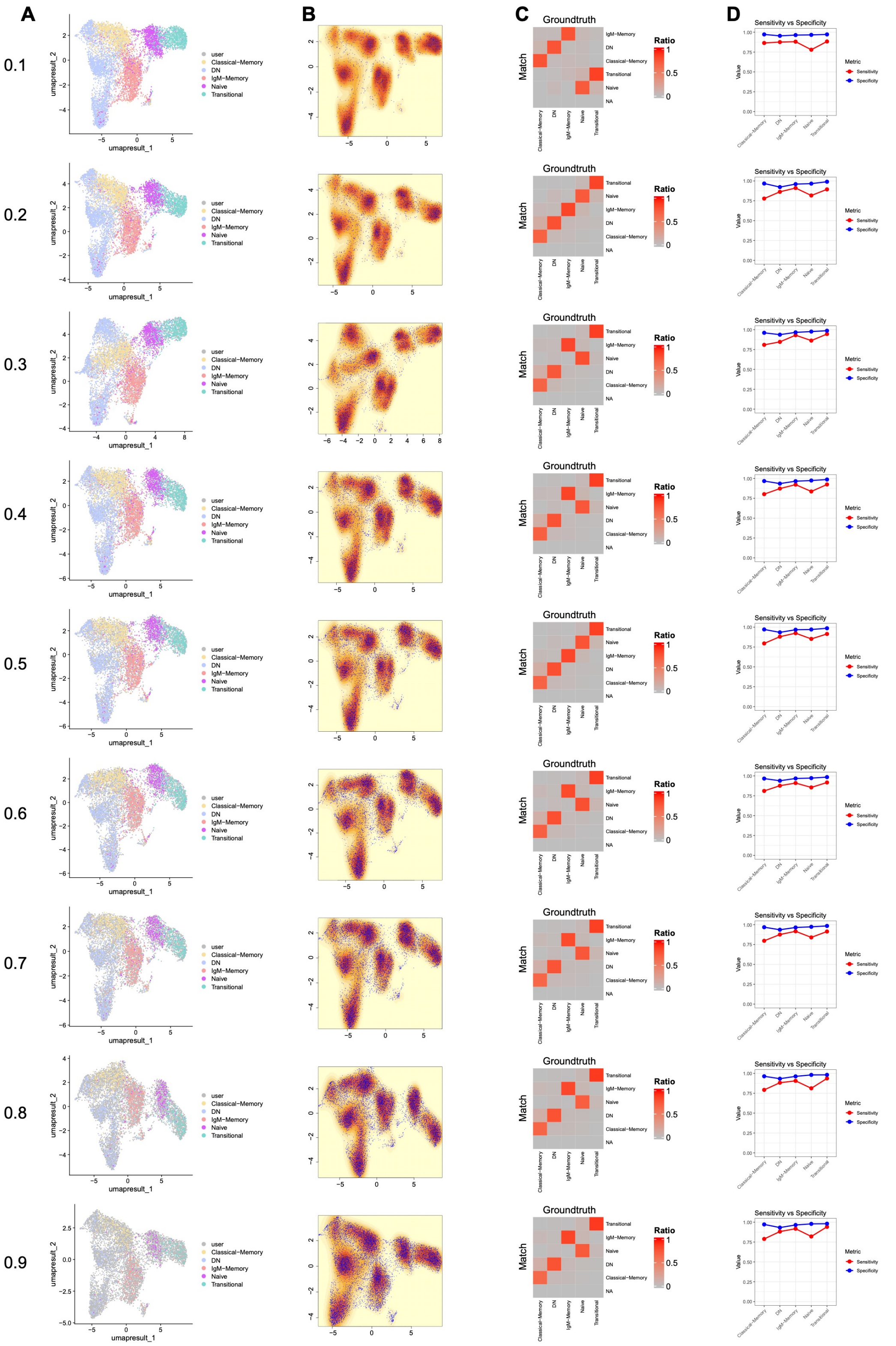


***Figure S2.*** *Results of STEVE using B cells annotated into five functional subtypes across proportions (0.1 to 0.9) of user data randomly sampled from the full dataset.*

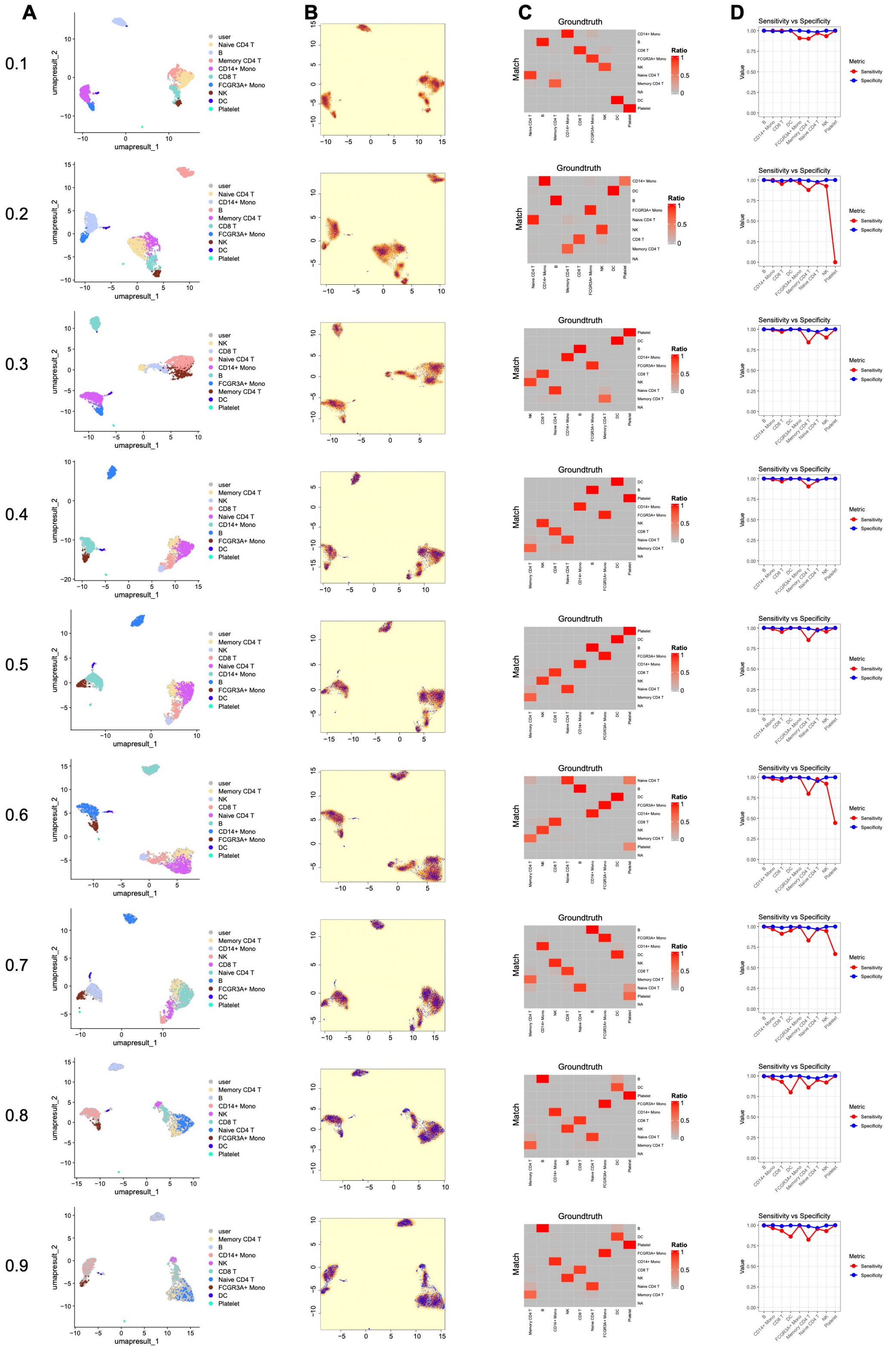

***Figure S3.*** *Results of STEVE using PBMCs from 10x Genomics across proportions (0.1 to 0.9) of user data randomly sampled from the full dataset.*

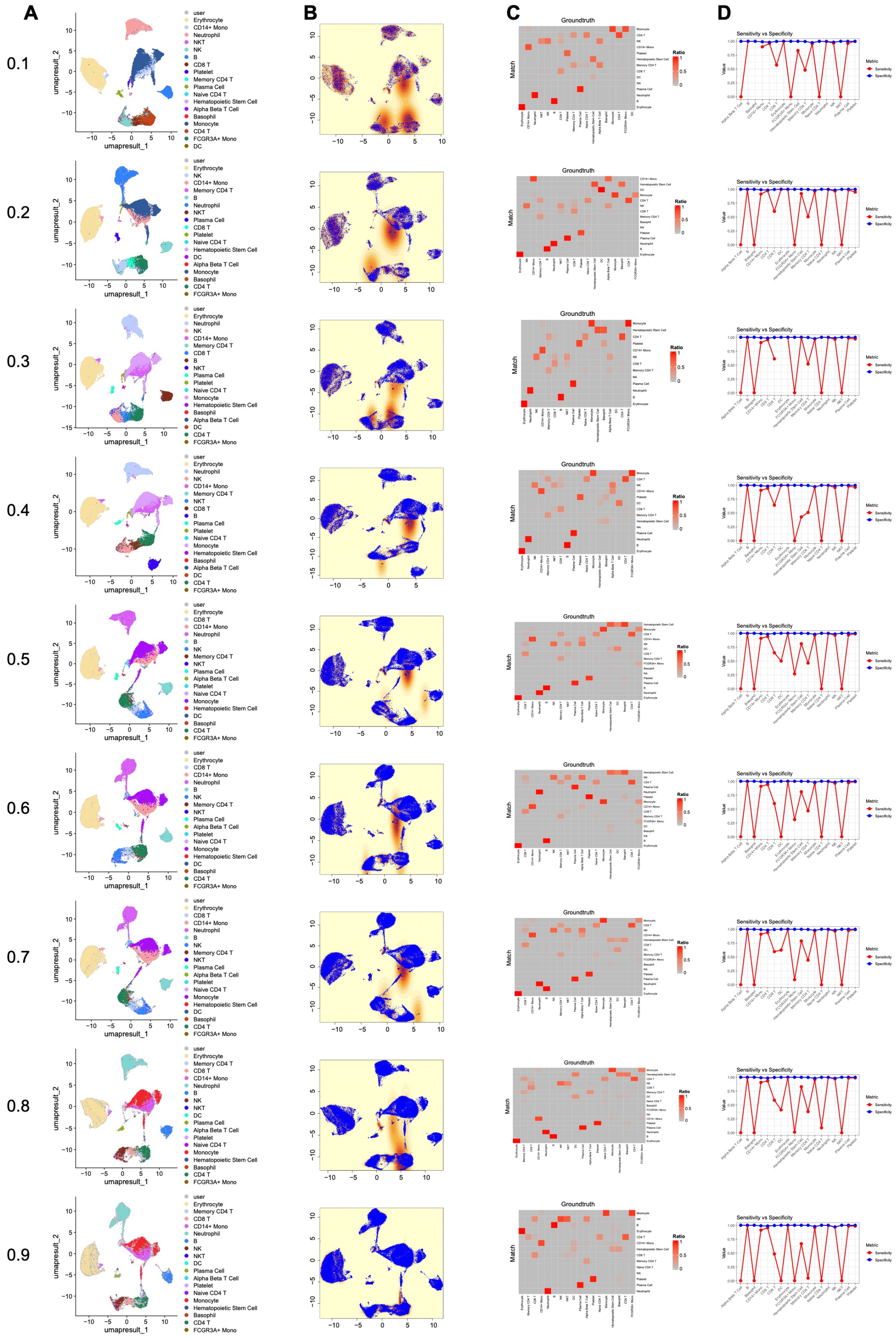

***Figure S4.*** *Results of STEVE using blood cell annotations from the Tabula Sapiens project across proportions (0.1 to 0.9) of user data randomly sampled from the full dataset.*

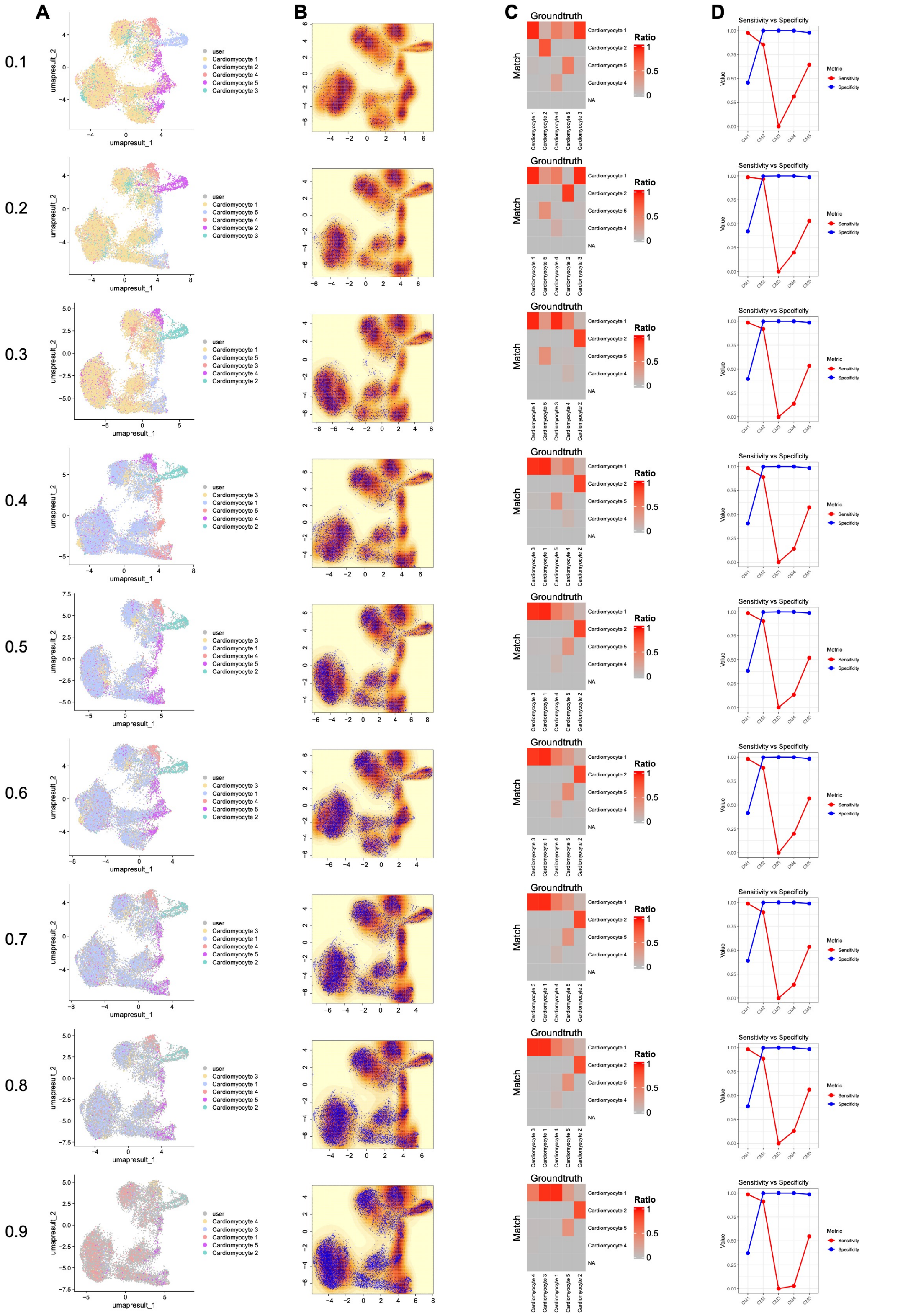

***Figure S5.*** *Results of STEVE using cardiomyocytes annotated into five functional subtypes across proportions (0.1 to 0.9) of user data randomly sampled from the full dataset.*

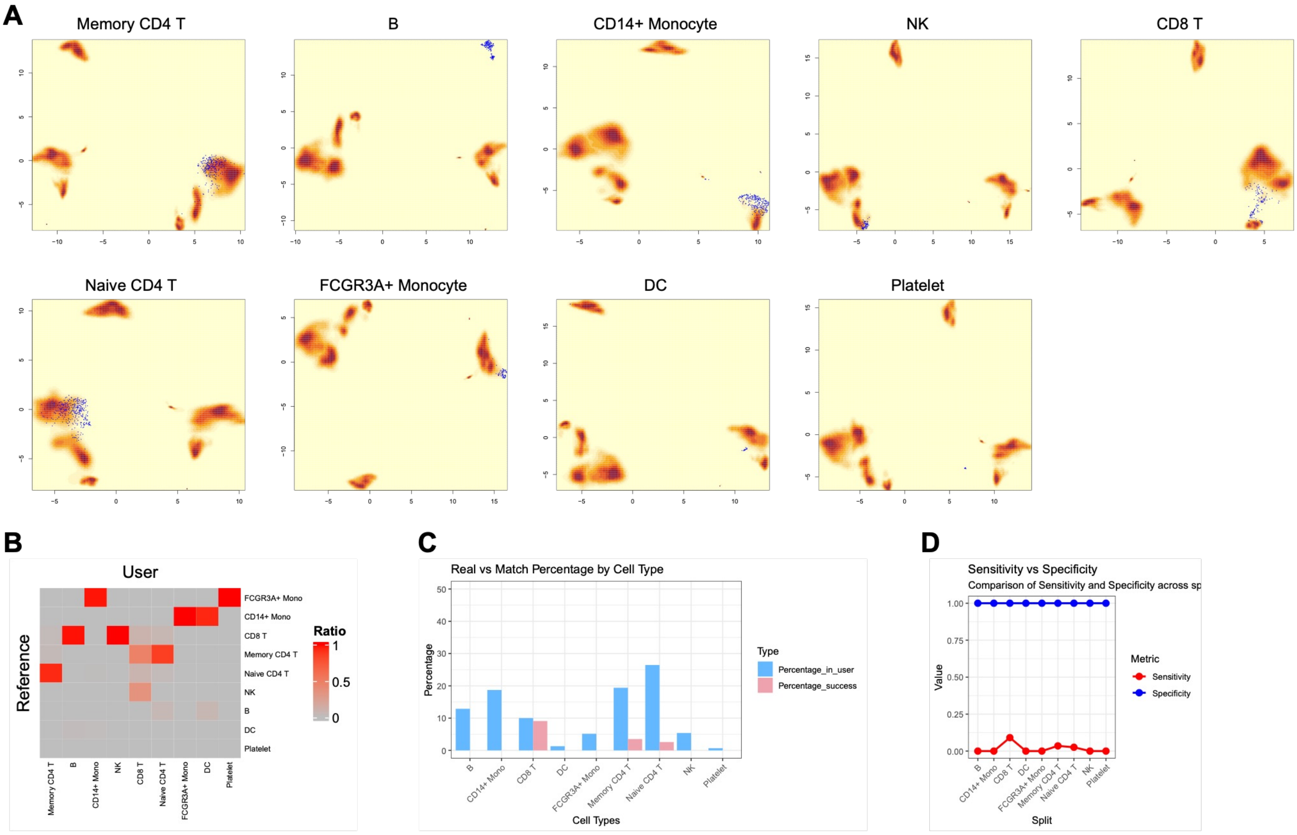


***Figure S6. Novel Cell Evaluation (10x Genomics PBMCs).***

*Here, we follow the same pattern as Figure3A-D, but for the 10x Genomics PBMCs dataset.*

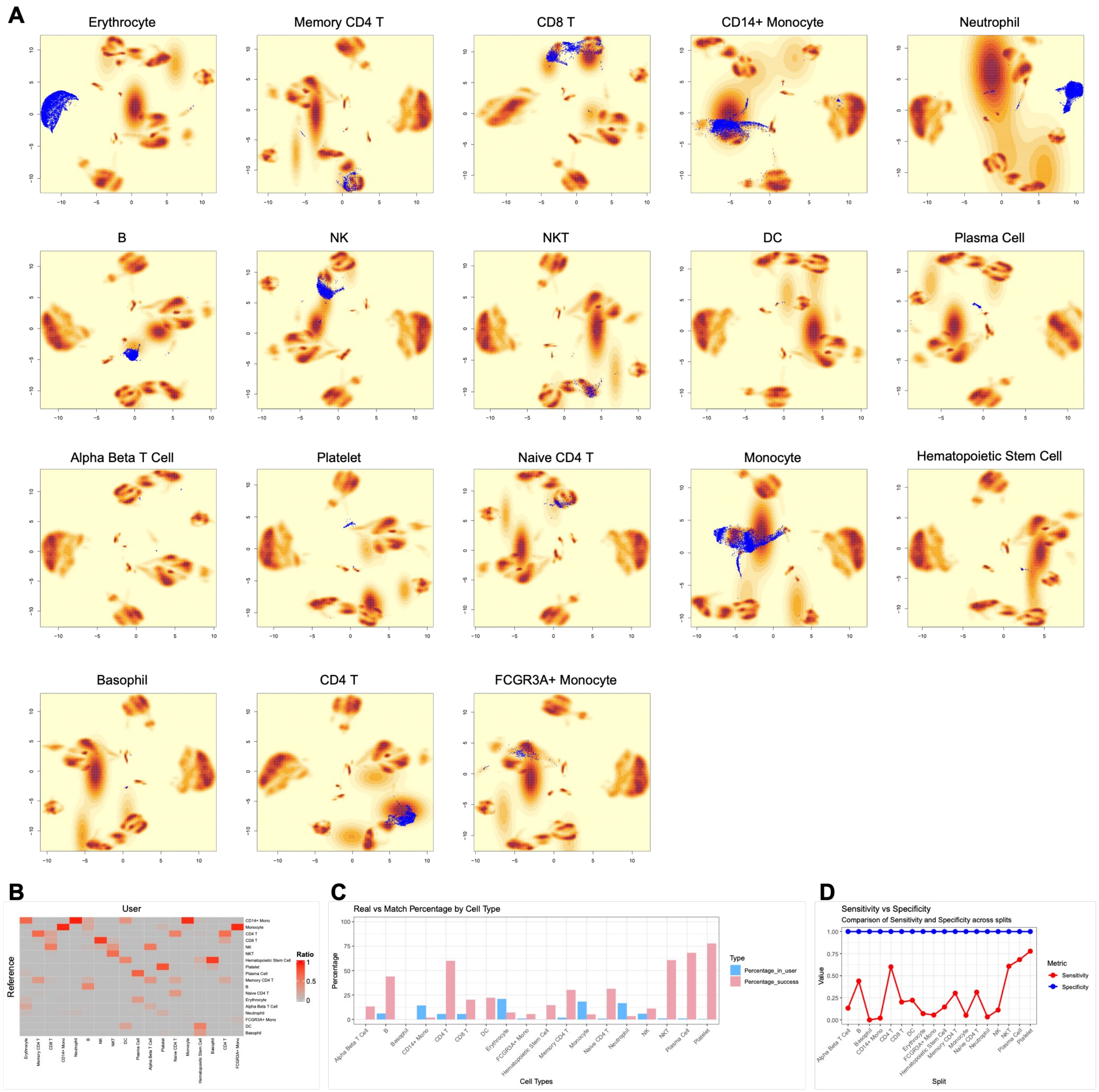


***Figure S7. Novel Cell Evaluation (Tabula Sapiens).***

*Here, we follow the same pattern as Figure3A-D, but for the Tabula Sapiens dataset.*
